# Effects of color-enhancing filters on color salience in normal trichromats

**DOI:** 10.1101/2025.10.14.682093

**Authors:** Camilla Simoncelli, Don McPherson, Kenneth Knoblauch, Michael Webster

## Abstract

Notch filters can alter color contrasts by selectively filtering different spectral bands of the stimulus and have been developed to enhance reddish-greenish contrasts for color-deficient observers with anomalous trichromacy. We examined the effects of such filters on color salience for normal trichromats, using a visual search paradigm where the task was to locate a color target superimposed on a variegated chromatic background, similar to foraging for fruits among foliage. Background colors varied along a bluish-yellowish or purpliish to yellow-green (short-wave cone isolating) axis, roughly spanning the range of dominant color variations in arid or lush environments. Target colors sampled a wide range of hue angles and contrasts. Testing was conducted on a computer monitor, with the filter effects simulated by calculating corresponding chromaticities with or without the filter for naturalistic (Munsell) reflectances. The filter evaluated (Enchroma SuperX® glasses) was designed to increase color contrast along a magenta-green axis. Consistent with this, search times for targets on the blue-yellow background were significantly faster for the filter condition, because the filter increased the target-background color difference. Alternatively, overall differences in search times were not observed for the S-cone background. The differences on the two backgrounds could be qualitatively accounted for by the relative salience of the stimuli predicted by a perceptual color space (CIELAB). Our results demonstrate the efficacy of the filters for enhancing visual performance for normal trichromats and naturalistic tasks, and illustrate how these effects depend on the potential color characteristics of the environment.

## Introduction

Normal human color vision depends on sampling the light spectrum by three types of cone photoreceptors maximally sensitive to short (S), medium (M), or long (L) wavelengths in the visible spectrum. Information about the spectral content of the light is contained in the differences or relative responses of the different cones, manifested in the underlying cone and color opponency in post-receptoral color processing. In a fixed context, larger differences correspond to stronger color contrasts or roughly to more saturated colors, while the type of difference (e.g. which cone is stimulated more) corresponds to the chromatic direction or roughly to hue.

The spectrum of a visual stimulus depends on both the characteristics of the illumination and the surface materials (Foster, 2011). Color contrasts in a scene could be enhanced by varying the spectrum emitted by the illuminant and is the principle employed in wide color gamut lighting and displays, which use spectrally narrow primaries to increase the range of spectral differences (Feng et al., 2016; Ilic et al., 2022; Wen at al., 2021; Zhu et al., 2015;). Alternatively, the spectra can also be shaped by filtering the light reaching the eye. Notch filters block or transmit different portions of the visible spectrum, and have been developed to increase the relative differences in the signals of the L and M cones as an aid for anomalous trichromacy (the most common form of color deficiency, resulting from a reduced separation in the peaks of the two longer-wavelength sensitive cones) (Somers & Bosten, 2024). A number of recent studies have evaluated the efficacy of the filters for color perception and performance for individuals with color deficiencies, and have documented conditions under which the filters do or do not have significant impact (Gómez-Robledo et al., 2018; Lynch & Ng, 2024; Marques et al., 2023; Moreland et al., 2022; Nascimento & Foster, 2022; Patterson et al., 2022; Pattie et al., 2022; Rabin et al., 2022; Somers & Bosten, 2024; Somers et al., 2024; Werner et al., 2020).

Unlike these previous studies, in the present work we instead examine the potential impact of spectral notch filters on *normal* color vision, using filters designed for normal trichromatic vision and testing observers with normal trichromacy. We also focused on how the changes in color contrast could impact the salience or conspicuousness of suprathreshold colors, rather than on threshold discrimination. Color vision above threshold manifests different characteristics of spatial and chromatic selectivity (e.g. Shapley et al. 2025), is altered by potential effects of response nonlinearities and gain (e.g. Knoblauch et al. 2020; Robinson et al. 2023; Tregillus et al. 2021), and is arguably more relevant to many ecologically important judgments. To assess color salience over a wide range of chromatic contrasts, we used a color search task in which observers had to locate a color target embedded in a color-varying background. Visual search is a ubiquitous natural behavior central to many visual tasks (Wolfe, 2020). The properties and characteristics of visual search have been studied extensively, and include many cognitive (e.g., goals) and sensory (e.g., stimulus salience) factors (Wolfe & Horowitz, 2017). Color provides important information for visual search (D’Zmura, 1991; D’Zmura & Mangalick, 1994), and this may have been one of the forces driving the evolution of color vision in many species. For example, a prominent theory of primate trichromacy is that it evolved to help find ripe fruit among foliage (Osorio & Vorobyev, 1996; Regan et al., 2001).

To measure color search, we used a “color foraging” paradigm developed by McDermott et al. (McDermott et al., 2010) that simulates search for colored “fruit” on densely colored backgrounds, and examined how performance on the task was affected by the changes in chromatic contrast introduced by a notch filter. The specific filter we tested was the Enchroma SuperX® glasses (EnChroma Inc., Berkeley, California, USA), which were designed for normal trichromats, and which amplify chromatic contrasts or colorimetric purity along a roughly magenta-lime axis. As illustrated below, the filtering can selectively increase the color contrast of some broadband spectra by up to 50%. Previous studies of color search have shown that search improves as the difference between the target color and background color (or color axis) increases (Bauer et al., 1996; D’Zmura, 1991; Manalansan et al., 2025; McDermott et al., 2010; Nagy & Sanchez, 1990). We therefore expected that the effects of the filters on the search task would be consistent with the chromatic differences they introduced in the stimuli, and with how these depend on the specific chromatic properties of the targets and backgrounds.

## Method

### Participants

Ten color-normal observers (mean age ± SD = 27.5 ± 4.4) were tested in the Visual Perception Laboratory at the University of Nevada, Reno (UNR). Most were undergraduate or graduate students at the university, who received course credit for participation. All had normal or corrected-to-normal vision and normal color vision as assessed by a battery of screening tests including the Ishihara, HRR and Dvorine plate tests, and the Farnsworth-Munsell 100 hue Test. Participation required written informed consent, and all procedures were in accordance with the Code of Ethics of the World Medical Association (Declaration of Helsinki) and approved by the Institutional Review board of the University of Nevada, Reno. A single session of the experiment lasted 1-1.5 hours, depending on the participant.

### Stimuli

All stimuli were presented on a 32’’ Cambridge Research Systems Display++ controlled by a computer running experimental code written in Matlab and PsychToolBox. The monitor primaries were calibrated with a spectroradiometer (Photo Research PR 655). The monitor was driven at a refresh rate of 120Hz and had a resolution of 1920 x 1080 pixels.

Chromaticities were defined within a version of Derrington-Krauskopf-Lennie cone-opponent color space, which represents the chromatic plane in terms of axes corresponding to variations in the LvsM cone and the SvsLM cone-antagonistic signals at constant luminance (L+M) (Derrington et al., 1984). The cone spectra were based on the Stockman-Sharpe fundamentals (Stockman & Sharpe, 2000) and the space was defined so that the gray origin corresponded to the chromaticity of illuminant D65 and a luminance of 20 cd/m^2^. Units along the two axes were scaled so that a distance of 1 unit corresponded roughly to the threshold for detecting a chromatic change away from white along each axis, based on measurements from prior studies, and so that positive values along the SvsLM axis corresponded to +S increments. The equiluminant plane in the resulting space is related to the MacLeod-Boynton chromaticity diagram by the following equations:

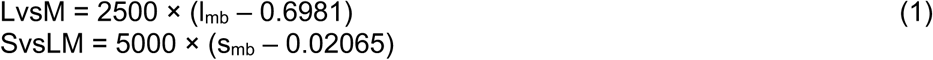

where 0.6981 and 0.02065 represent the MacLeod-Boynton L/L+M) and S/(L+M) values of the D65 gray point based on the Stockman-Sharpe 2-deg fundamentals (Stockman, 2019). These fundamentals represent the standard observer at the center of gaze, and thus do not account for changes in sensitivity with retinal eccentricity or for individual differences across observers (Rider and Stockman, 2025), though these normal variations would not introduce qualitative changes in the predicted effects of the filters on the cone responses.

The use of a monitor was necessary to allow systematic variation and control of the stimulus and experiment. However, for our study we were interested in the impact of the notch filters on naturalistic spectra, and not on the spectra generated by a typical RGB display. Monitor spectra are relatively narrowband and in particular already have lower emission at wavelengths that the notch filters are designed to block. Thus, the filters have more limited effects on colors viewed on a monitor (Somers and Bosten, 2024). To overcome this limitation, testing was performed without wearing the filters, and instead we simulated the filter effects by modeling their impact on naturalistic reflectance spectra and then presenting the monitor stimuli with the same chromaticity. The natural spectra were based on the first 3 principal components of variation in the Munsell reflectance spectra (Cohen, 1964) when shown under D65 illumination. This transform allowed us to first specify the DKL coordinates of our stimuli and then construct the corresponding Munsell spectra (Ilic et al., 2022; Webster & Mollon, 1995). These simulated spectra were then filtered by the transmittance of the notch filter and weighted by the LMS cones to yield the predicted cone responses. Filtering necessarily reduces the average luminance of the stimuli and potentially also changes the average chromaticity. To control for these differences, the predicted cone responses to the stimuli were rescaled (as in the standard von Kries transform) so that the mean background level after filtering equaled the response without filtering. This step maintained all backgrounds at the same mean luminance (20 cd/m^2^) and chromaticity (of D65), and was equivalent to assuming complete chromatic adaptation to the background.

The panels of Figure 1 illustrate the DKL chromatic plane (left) and the transmittance of the SuperX filter (right). For the naturalistic spectra we modeled, the effect of the filtering is to increase chromatic contrast roughly along the reddish-cyan LvsM (0-180°) axis of the space, with the least chromatic change along a bluish to yellowish-orange axis (roughly 135-315°) (Figure 2).

**Figure 1.**
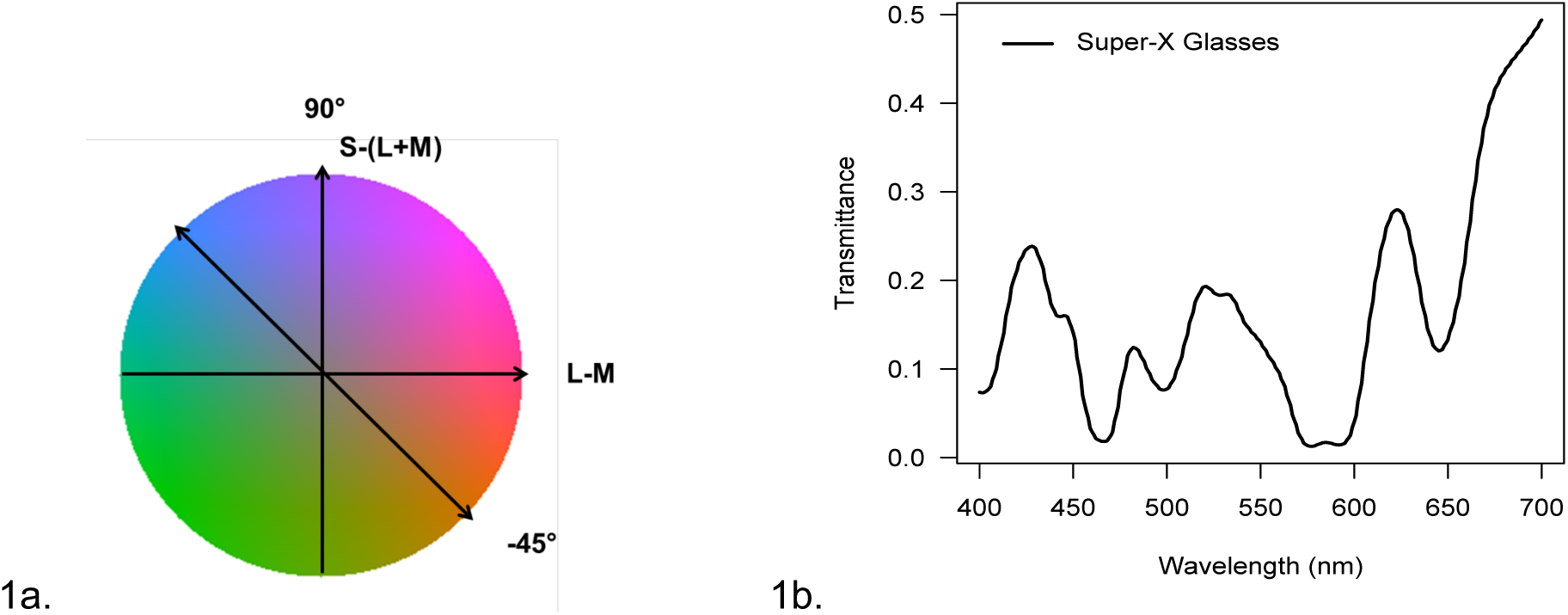
(a) 90-270° and 135-325° (shown and referred to as 90° and –45°) background chromatic axes in the DKL color space. (b) Enchroma SuperX® glasses spectra transmittance (400 – 700 nm).

**Figure 2.**
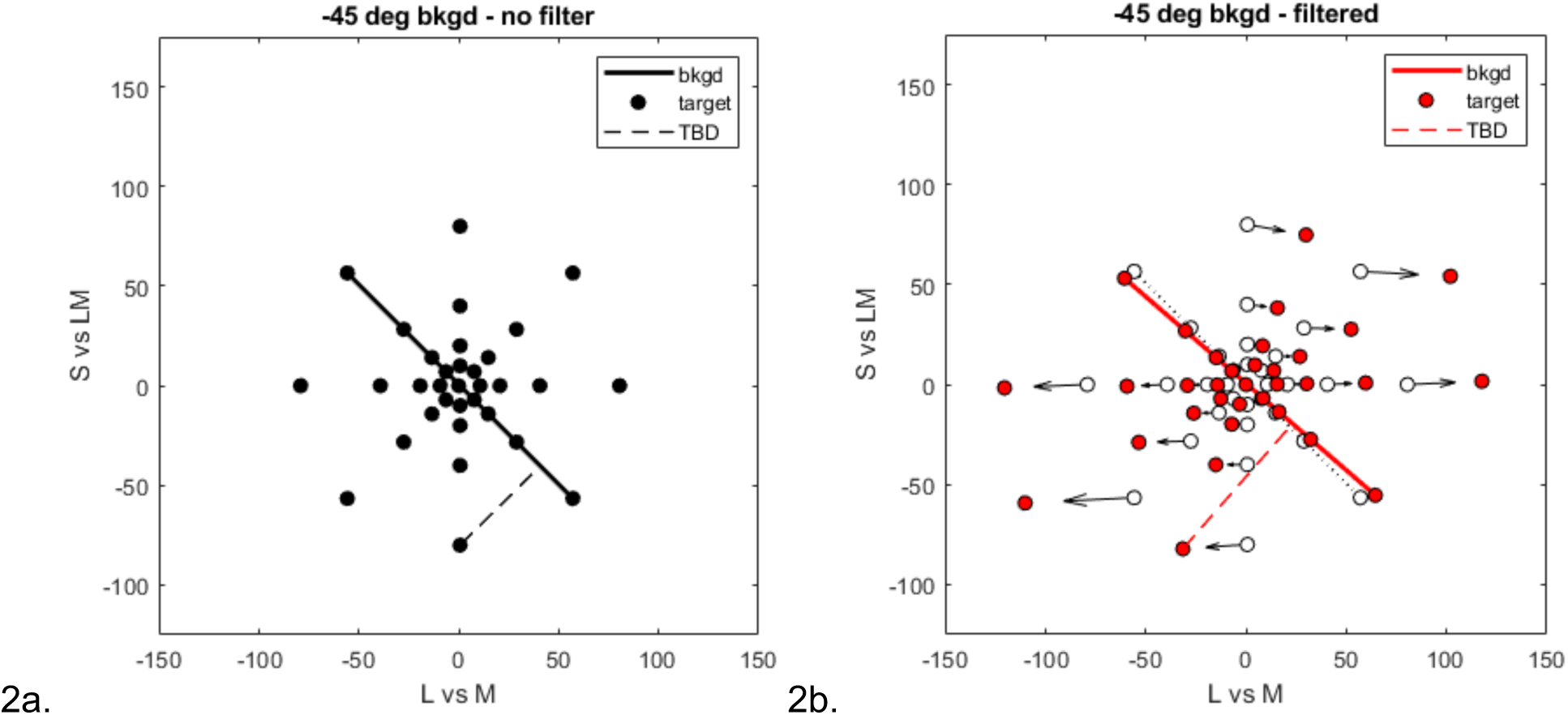
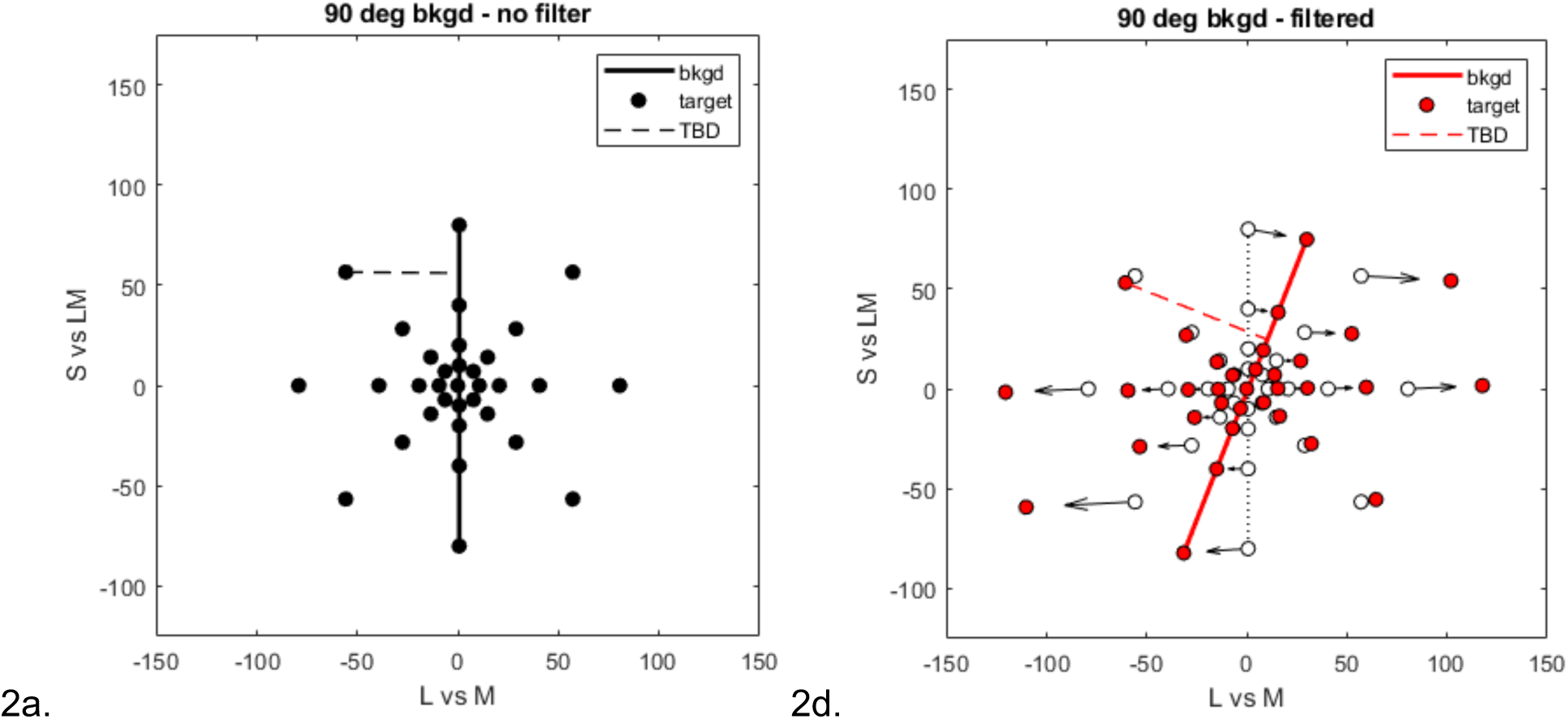
Top: targets and background coordinates in the LvsM and SvsLM plane without (a) and with (b) the filter for the –45° background. Bottom: targets and background coordinates without (c) and with (d) the filter in the 90° background. Note that the same set of targets were used for both backgrounds, so that only the chromatic axis of the background differs between the upper and lower panels.

For the experimental stimuli, the background colors were varied along two different axes in the plane, chosen to span the range of variation in the principal axes of color gamuts in typical natural scenes (Ruderman et al., 1998; Skelton et al., 2024; Sumner and Mollon, 2000; Webster et al., 2007; Webster & Mollon, 1997).. For one, the colors varied along angles of 135-315° axis, again a bluish to yellowish-orange direction typical of arid and panoramic scenes. This axis is thus similar but not identical to the unique blue-yellow axis of classic perceptual opponency. For the second axis, the background colors were confined to variations along the 90-270° SvsLM axis, which varies from a purple to yellow-green. This axis corresponds to the tritan line along which signals only in S cones vary at constant luminance, and is more characteristic of scenes dominated by lush foliage. For shorthand, we refer to these two conditions as the –45° and 90° axes respectively. As discussed below, the different axes also allowed us to assess how the impact of the filter on color salience depended on the background colors. For each background, the chromaticities of individual elements were selected randomly from contrasts spanning +/-80 units. The luminance of each background element was also randomly varied by +/-30% relative to the 20 cd/m^2^ mean of the background, in order to mask luminance differences as a potential cue to the target (Regan et al., 1994). Target colors were instead chosen to sample 8 different color directions or hue angles, (at steps of 45° around the cone-opponent plane, Figure 1a) each shown at 4 different contrast levels ranging from 10 to 80 units in 0.3 log steps, and all at the mean luminance before the filtering. The filtering also changed the relative luminances of the target and background colors, and this effect was included in the rendered stimuli. However, these luminance changes were less than +/-6% and thus small relative to the chromatic changes and to the random luminance variation added to the background.

Figure 2 shows the chromatic coordinates of the targets and the two backgrounds. Note that the same targets were used for both backgrounds. Note also that the targets include colors at different angles relative to each background and include colors along the background axis. The angles and contrasts of both the targets and backgrounds are altered by the filters. In the analyses below, we represent the effective contrast of the target by the distance of the target chromaticity to the background (TBD) axis. This varies from 0 for all targets along the background axis to the full nominal contrast for the targets at angles perpendicular to the axis.2

To display the stimuli, a variegated color background was displayed composed of 10,000 superimposed ellipses whose minor axis randomly varied from 0.2° to 0.39° of visual angle and major axis from 0.62° to 0.8°, according to a uniform distribution. The ellipses’ positions and orientations were also randomly varied. The ellipses occluded those below them and the large number was to ensure the screen was fully tiled. The circular target had a diameter of 0.5° and was randomly positioned on top of the background, so that it was always fully visible. The circular shape cue meant that the target could be detected even when the color was along the background axis, which allowed observers to identify the target on all trials. Examples of the backgrounds and targets are illustrated in Figure 3.

**Figure 3.**
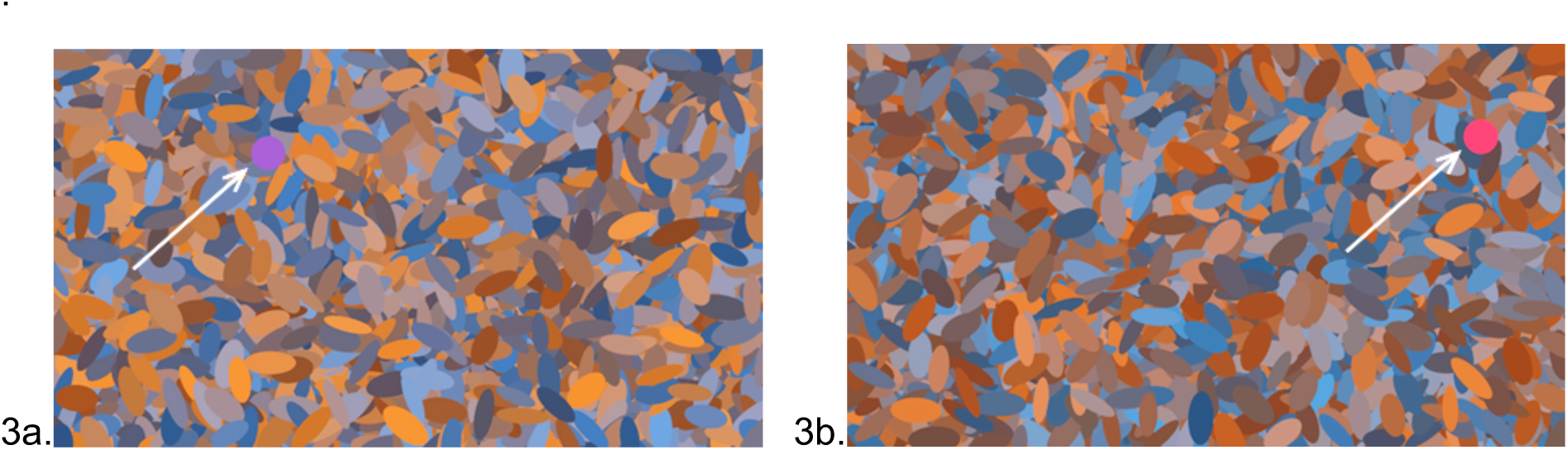
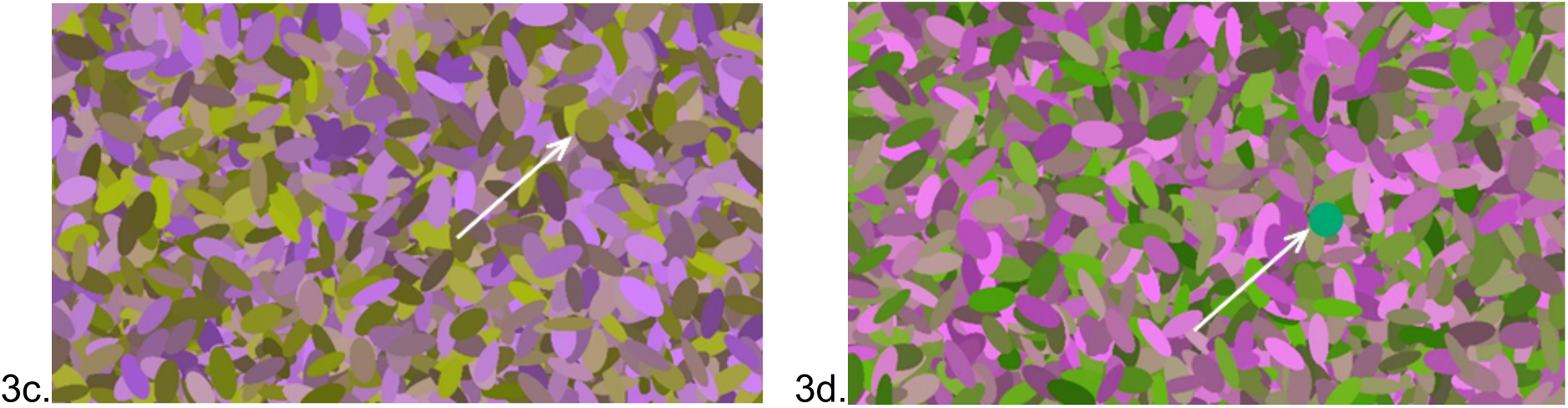
Examples of the stimuli for –45° background without (a) and with (b) the filter simulation, and for 90° background without (c) and with (d) the filter simulation. Arrows show the location of the targets, which vary in visibility based on the color difference from the background.

### Procedure

Observers viewed the display binocularly in an otherwise dark room and were seated at 120cm from the monitor, at which the full-screen background subtended 20° x 11.6°. There was not a chin rest or head restraint so the actual distance and visual angles are approximate. On each trial the target was displayed at a random location on the left or right side of the display, and the participant had to identify the side by using a handheld keypad. There was no time limit to respond, and a response was required in order to advance to the next stimulus. Responses were signaled by a tone but feedback for correct responses was not provided. Each response was followed by a random delay of 2-3 sec before the next stimulus, during which the screen remained the uniform gray.

Each participant completed 20 repetitions for each target/background condition (one experimental block). Each session was composed of four experimental blocks (two with the filters and two without the filters) with the order of the filter condition counterbalanced and unknown to participants. For each background two sessions were completed on separate days; the whole experiment, thus, lasted 4 days for the 4 sessions, with the time of a single session lasting between 1 and 1.5 hrs. The order of the two backgrounds was also counterbalanced between the participants. Results reported are for the aggregate data across observers: 2560 total observations per observer were recorded (1280 for each background).

## Statistical Analyses

To formally assess the results, the data were analyzed in the R environment for statistical computing (R Core Team, 2024). For both backgrounds, we conducted sign tests of the median reaction times to compare the filter vs. no-filter condition results, for session 1, for session 2 and for the two sessions together, both for between-observer and within-observer comparisons. We then fitted nonlinear mixed-effects models (NLME) to account for the effects of the Target-Background Difference (TBD). We initially also analyzed the Target Contrast, which showed a linear dependence on the dependent variable after it had been transformed logarithmically and similar filter effects to the TBD. A principal component analysis of the explanatory variables, however, indicated that the TBD and Target Contrast were highly correlated. The first principal component depended on both variables with approximately equal weighting. Therefore, we opted to analyze just the TBD here.

Histograms of reaction times (RT) are typically highly positively skewed, which leads to overestimated and biased confidence intervals using standard mean and variance estimators, and thereby leading to reduced power in hypothesis testing (Zandt, 2002). We evaluated the residuals of a linear model including all variables with Box-Cox power transformations (Box & Cox, 1964) using the *boxcox* function in the **MASS** package (Venables & Ripley, 2002). The analysis suggested that the reciprocal of the RT or speed would stabilize the variance and approximately normalize the distribution of responses, considered desirable properties of a good measurement scale (McCullagh, 2019). Simulations have indicated that the inverse transform minimizes the effects of outliers (Ratcliff, 1993), An example of the extreme skewness in the RT distribution for one observer for the 90° background condition, both filter conditions confounded, is shown in Fig 4a. The same data are shown in Fig 4b with the histogram now plotted as a function of the speed or reciprocal reaction time. As the results were similar for all subjects for both background conditions, the speed was analyzed as the dependent variable in subsequent analyses. The exception is the initial analysis of medians, which is a robust (but less powerful) statistic that depends much less on the distribution shape.

**Figure 4.**
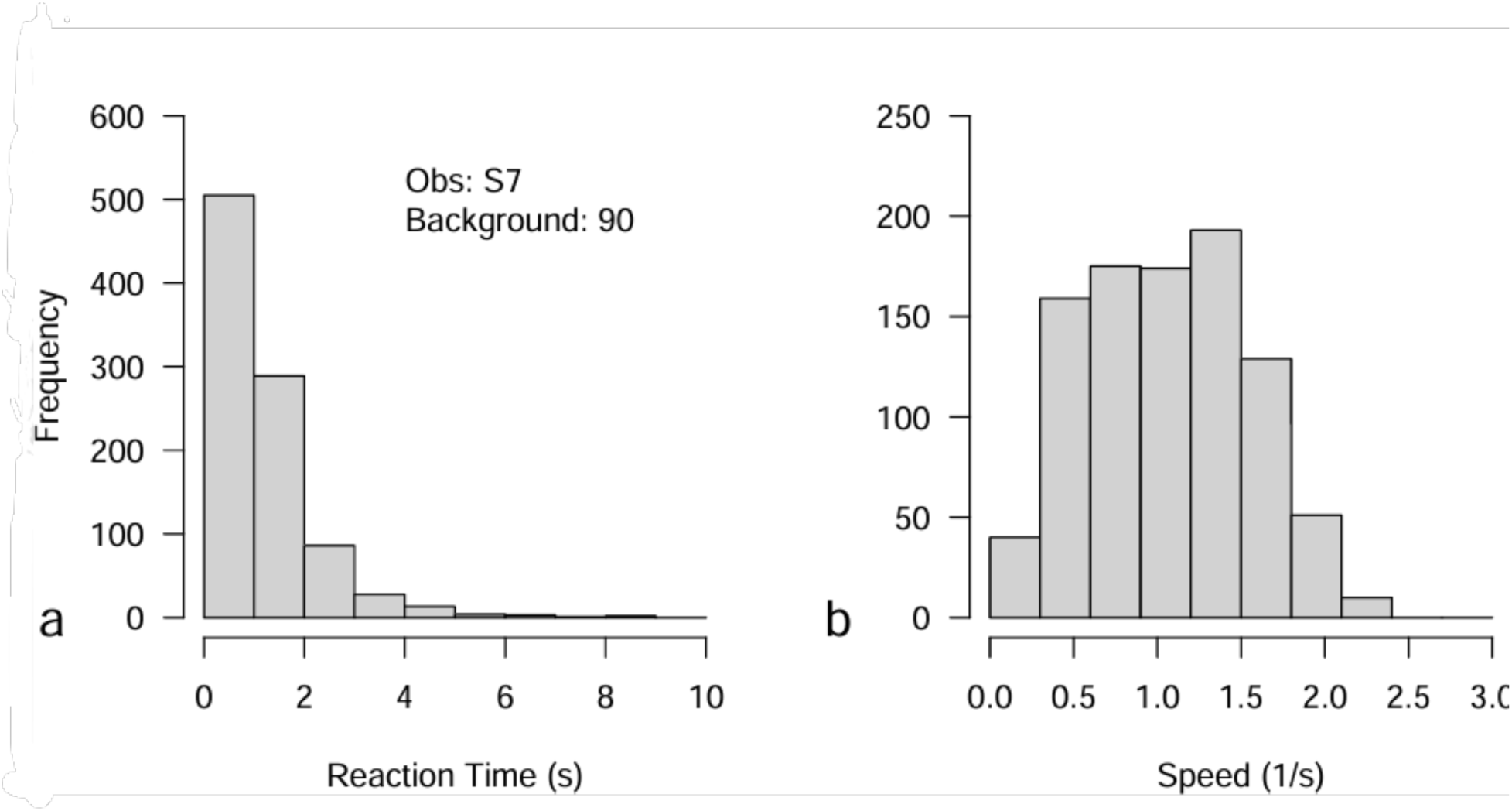
(a) Histogram of frequencies of reaction times in seconds for observer S7 for the 90° background condition. (b) Histogram of frequencies of reciprocal reaction times in inverse seconds for the same data in part a).

Nonlinear mixed-effects models were fit to the transformed data using the nlme function in the **nlme** package (Pinheiro et al., 2025; Pinheiro & Bates, 2000) to model the effect of the filter as a function of TBD and Target Angle. Initial evaluations indicated that the results did not depend on Target Angle and it was excluded from the main analyses. A Michaelis-Menten function was found to provide a good description of the dependence of the speed on TBD. In preliminary analyses a power law was found to provide a similar description. However, as the best fits with the Michaelis-Menten function yielded a smaller AIC, we used this model for subsequent analyses. Random effects and then fixed effects were evaluated with χ^2^ statistics obtained from nested likelihood ratio tests. The full NLME initially fit to the data is formalized as

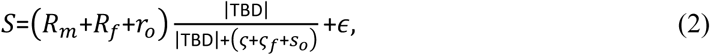

where

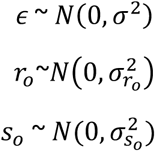

and where *S* is the speed (in reciprocal seconds), TBD the target-background distance (as defined below), the *R* and ς parameters indicating the maximum speed and TBD at 50% maximum response (semi-saturation constant) for the no filter condition, respectively. *R_f_* and ς*_f_* are the respective fixed effects of the filter on the two preceding parameters and *r*_0_ and *s*_0_ are random observer predictions of each of the preceding parameters, each assumed to be normally distributed with its own variance. The parameter ϵ is the residual random variation that is assumed to be normally distributed with mean zero and variance σ^2^. Thus, the full model has 3 variance terms and the correlation between the two parameter random effects to estimate in addition to the fixed effects of the no filter and change due to the filter on each of the parameters. Nested likelihood tests were performed initially to evaluate the significance of the random effect terms. Subsequently, the fixed effects of adding the filter were evaluated for each parameter, *R_f_* and ς*_f_*. We set the level of significance at α = 0.05 and report statistical analyses only for correct responses, which were overall very high across conditions and observers (99.1%). No outliers were removed.

## Results

Figure 5 shows the reaction times for detecting the targets under the 4 conditions tested: on the two different backgrounds, and with or without the filter. As illustrated in Figure 2, the RTs are plotted as a function of the TBD, which is defined by

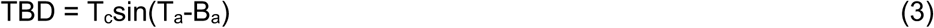

**Figure 5.**
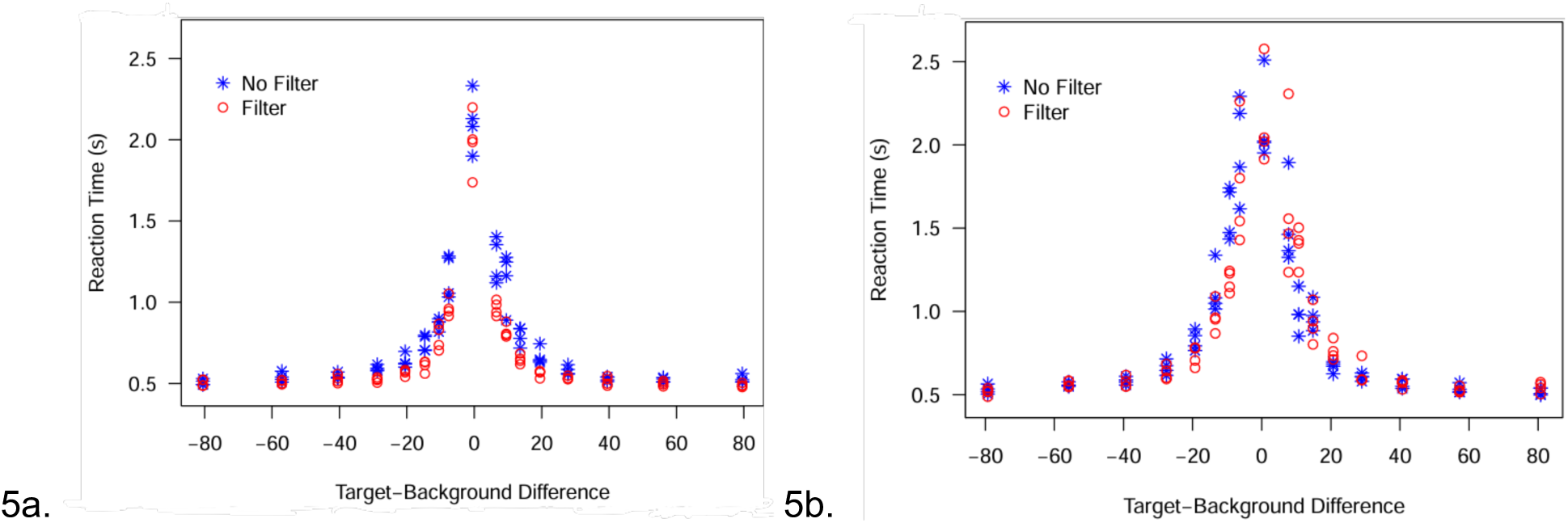
Reaction times plotted as a function of the target-to-background chromatic difference for the – 45° (a) and 90° (b) backgrounds. Blue asterisks represent the average reaction times in the no-filter condition, while red circles denote the corresponding values for the filter condition. Specifically, each data point reflects the mean across observers of the median reaction time calculated within each run, where the median is derived from five repeated reaction time measurements per observer. This approach ensures a robust central tendency measure by first summarizing individual trial variability within runs before aggregating across observers.

**Figure 6.**
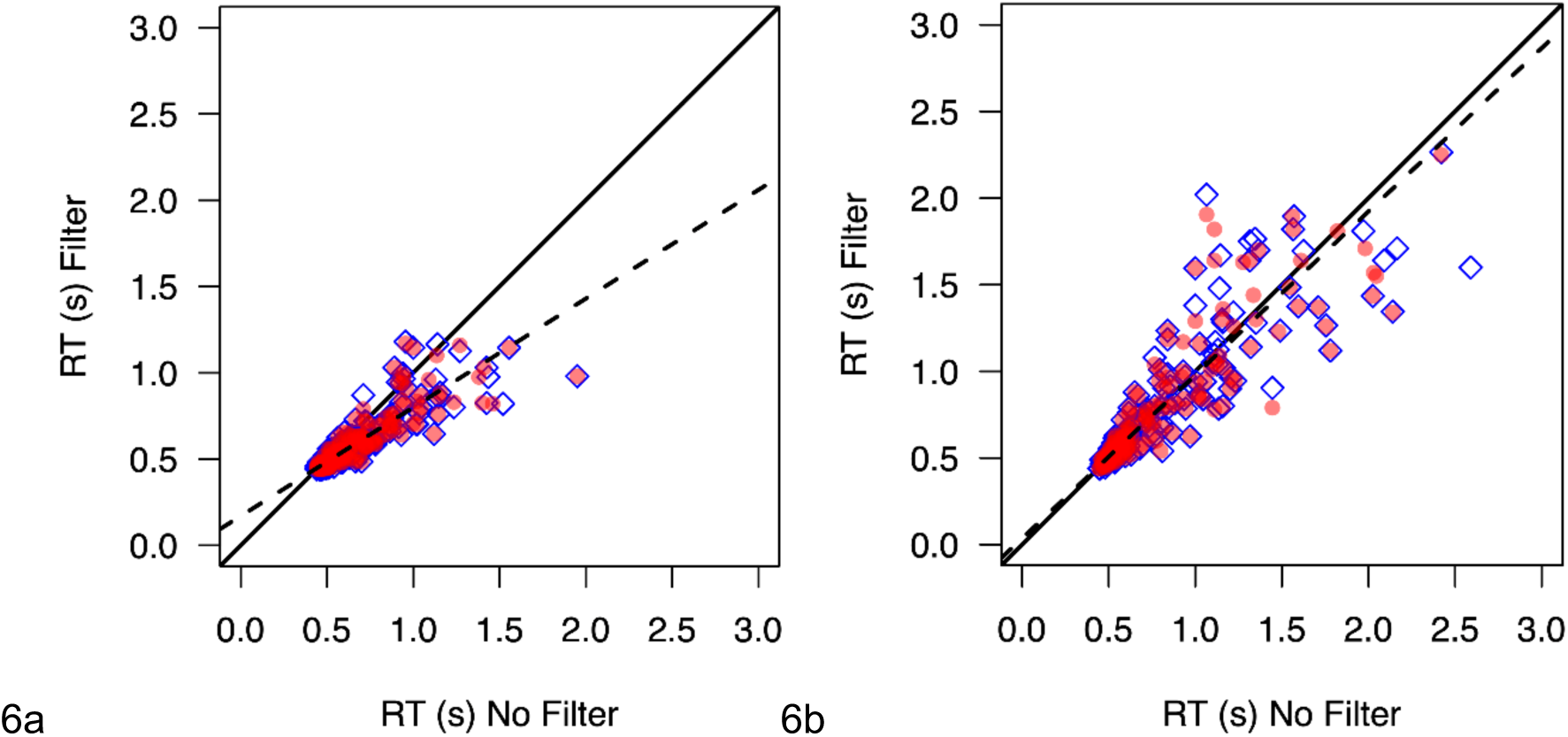
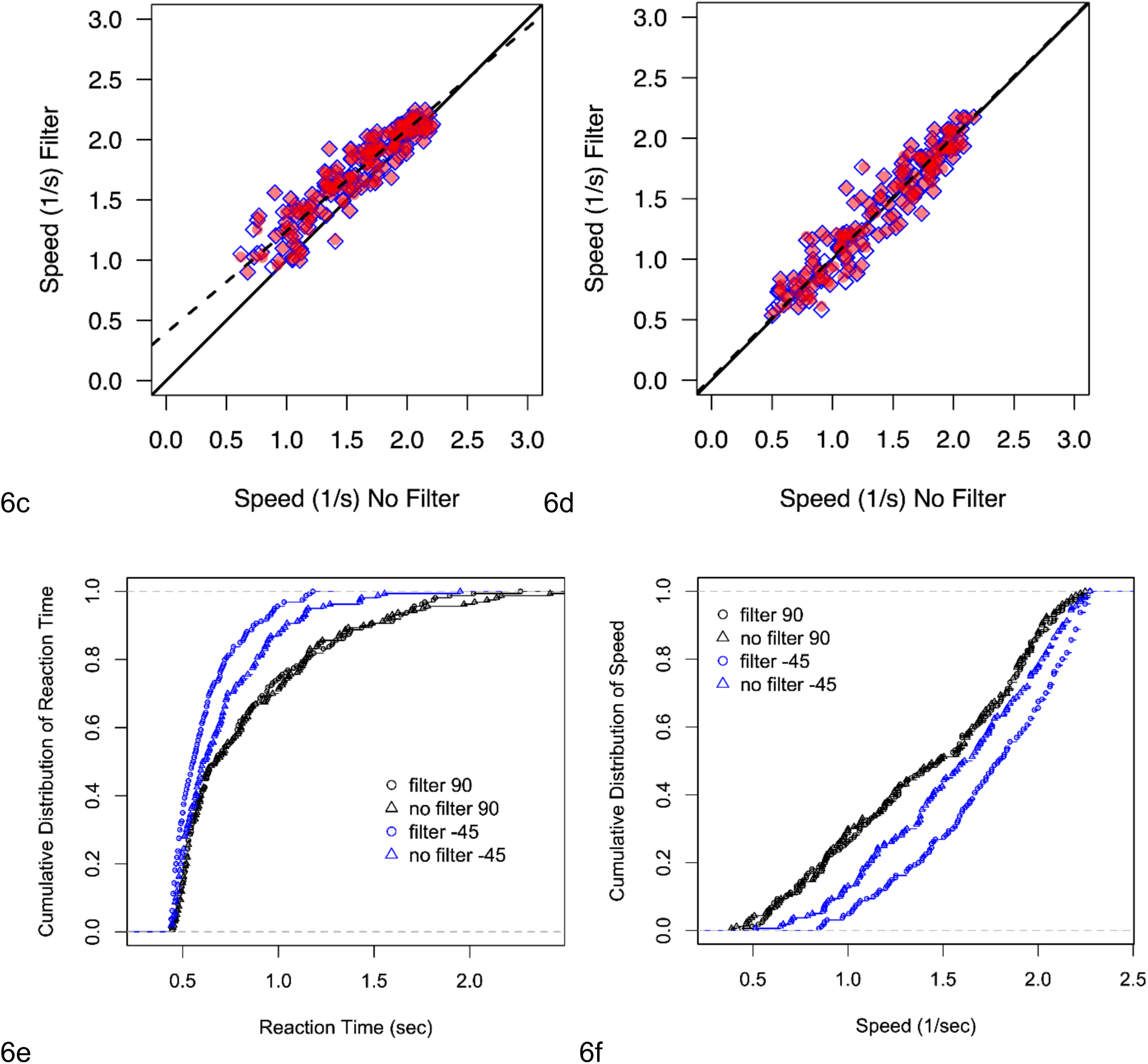
a and b) Average reaction times for the same targets on the –45 deg (a) or 90 deg (b) backgrounds compared for the filtered vs. unfiltered conditions. Symbols plot the RTs for all responses (blue diamonds) or correct responses only (red circles). Dashed line shows the best-fitting line. c and d) the same results after converting reaction time to speed. e and f) Cumulative probabilities of the reaction times (e) or speed (f) on the –45 (blue) or 90 deg (black) backgrounds.

where T_c_ equals the target contrast and T_a_ and B_a_ are the target and background chromatic angles, respectively. As in many previous studies, RTs decrease asymptotically as the separation between the target and background or distractor colors increases (Bauer et al., 1996; Manalansan et al., 2025; McDermott et al., 2010; McKeefry et al., 2003; Nagy & Sanchez, 1990; Vanston et al., 2021; Webster & Mollon, 1994). In our case, these effects are bounded by targets that “popped out” or were found rapidly when the color was very different from the background, to long search times when the target color fell within the background so that the targe could only be found by the shape difference. a (McDermott et al., 2010).

The questions of interest were whether the search times differed for the filter vs. no-filter condition, and whether this depended on the chromatic properties of the backgrounds and targets. To visualize these differences, in Figure the scatterplots directly compare the average reaction times (panels a and b) and speed (panels c and d) for each target under the filtered or unfiltered condition. For the –45° background, search times appeared consistently faster with the simulated filters. Alternatively, there is little evident difference for the 90° background. Note also that overall, the search times were substantially faster on the blue-yellow background than the SvsLM background. Finally, the effects of filtering and of the background axis are also more readily apparent when responses are instead plotted as the cumulative probability of the reaction times or speed for the different conditions (panels e and f). These patterns are confirmed statistically in the following analyses.

### -45° Background

Within-subjects comparisons for the –45° background revealed a global trend of the filter effect, without any distinction between the first and the second session: speed was higher in the filter condition. Sign tests for the between-subjects comparison showed statistically significant results across all the data: the filter had a greater effect both in the individual sessions (2-tailed: Session 1 – p = 0.002, Session 2 – p = 0.02) and in the two sessions together (2-tailed: p = 0.002). The assessment of the individual sessions was done to confirm that the differences were stable across the sessions. Participants were, thus, faster in the experimental blocks with the filters. The average filter vs. no filter difference for correct trials in session 1 was –207 ms, – 134 ms in session 2, and –171 ms in the two sessions together. The general accuracy was 99.4%, indicating that global performance was highly accurate.

Fitting the data to the NLME of Equation 2 proceeded in two steps. First, the random effects were evaluated to simplify them, if necessary, and then the fixed effects. Tests of the random effects indicated that the correlation between the two random effects was not significant (LRT: χ^2^(1)= 0.001, p = 0.97) permitting subsequent analyses to be performed with a diagonal covariance matrix. Random variances associated with the maximum value, 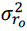 and the semi-saturation constant, 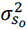, however, were required as leaving either of them out yielded a significant difference from the model containing both (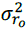, LRT: χ^2^(1)= 334.4, p << 0.001; 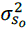, χ^2^(1)= 8.9, p = 0.003).

With the random structure of the model set, the fixed effects were evaluated. A filter effect on the maximum value parameter was not significant (LRT: χ^2^(1) = 0.339, p = 0.56), but the filter effect on the semi-saturation constant was (LRT: χ^2^(1) = 250.0, p << 0.001). A summary of the fixed and random effects parameter estimates are given in column 2 of Table 1 with their 95% confidence intervals in column 3.

**Table 1.** Estimates of fitted parameters and standard errors (SE) from best fitting NLME models for the – 45° and 90° deg data sets.

|  | -45 |  | 90 |  |
| --- | --- | --- | --- | --- |
|  | Estimate | 95% Conf. Int. | Estimate | 95% Conf. Int. |
| $R_m$ | 2.212 | (2.072, 2.351) | 2.336 | (2.183, 2.488) |
| $\zeta$ | 8.996 | (8.480, 9.512) | 15.303 | (14.346, 16.259) |
| $\zeta_f$ | -3.003 | (-3.305, -2.700) | -0.710 | (-1.220, -0.201) |
| $\sigma_{r_o}^2$ | 0.049 | (0.020, 0.119) | 0.057 | (0.023, 0.141) |
| $\sigma_{s_o}^2$ | 0.329 | (0.089, 1.216) | 1.184 | (0.814, 1.723) |
| $\sigma^2$ | 0.168 | (0.163, 0.173) | 0.181 | (0.177, 0.187) |

Figure 7a shows the average speed as a function of TBD for no filter (blue) and filter (orange) conditions. The error bars indicate 95% confidence intervals. The curves are based on the estimated fixed effects parameters from the best fit model for the no filter (solid) and filter (dashed) conditions based on the fixed effect parameters in Table 1, and indicate the generally faster responses to the filter condition. Note the similarity in the size of the confidence intervals across TBD, supporting the transformation of RT to speed to stabilize the variance. (Compare with the larger variation in RTs as a function of TBD in Figures 5). Concurrently, the filter had an approximately constant effect on all targets, independently of their angle, showing faster detection on experimental blocks with the filter on average (Figure 7b).

**Figure 7.**
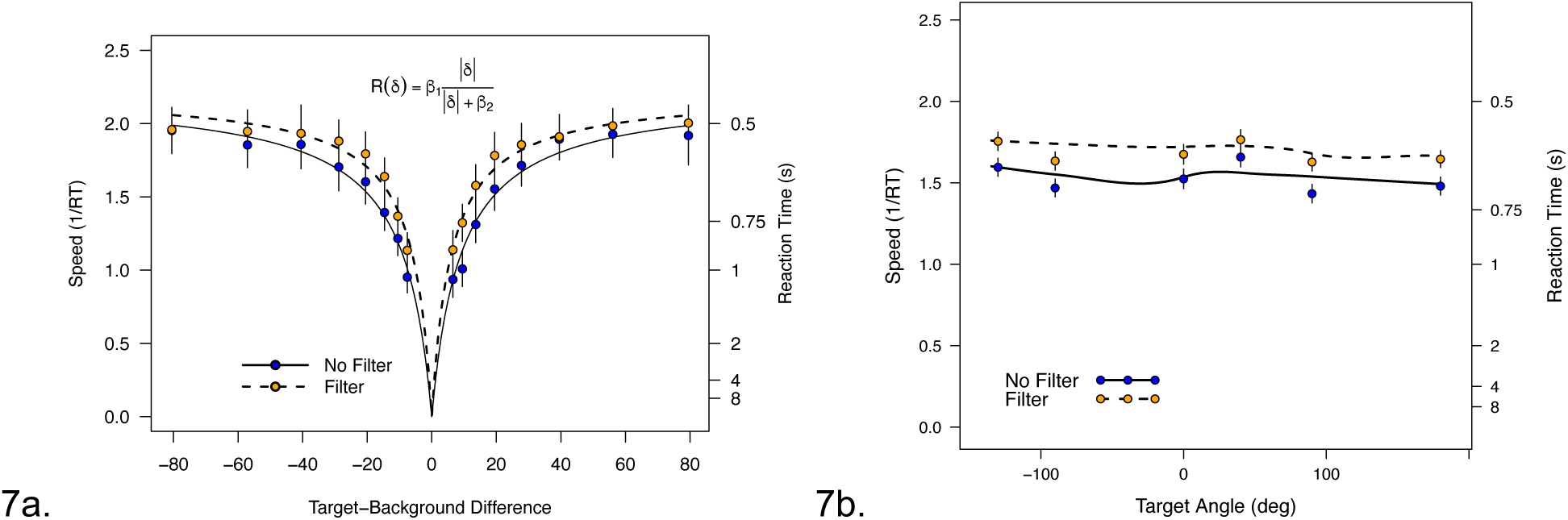
(a) Average speed (1/RT) with and without the filter as a function of the TBD. The curves indicate the fixed-effect model fit using the Nonlinear Mixed-Effect Model. Reaction times are indicated on the right ordinate. The Michaelis-Menten equation for the curves is indicated as an inset. Error bars are 95% confidence intervals. (b) Average speed as a function of Target Angle with the corresponding Reaction Times (in s) indicated on the right ordinate. The curves are local regression curves fit to the data. Error bars are 95% confidence intervals.

### 90° Background

Sign tests for the comparison between the filter and no-filter runs were not statistically significant for either individual session or the two sessions together (2-tailed: p=0.75, in all 3 cases). The average difference in session 1 was 81.4 ms, –33.2 ms in session 2, and 24.1 ms in the two sessions together. The global accuracy was 98.8%, similar to the –45° condition.

As with the –45*°* background, speed again varied systematically as a function of TBD with speed increasing with the magnitude of TBD (Fig 8a) and again was relatively independent of Target Angle (Fig. 8b). The differences between the no filter and filter conditions appear less obvious on this background.

**Figure 8.**
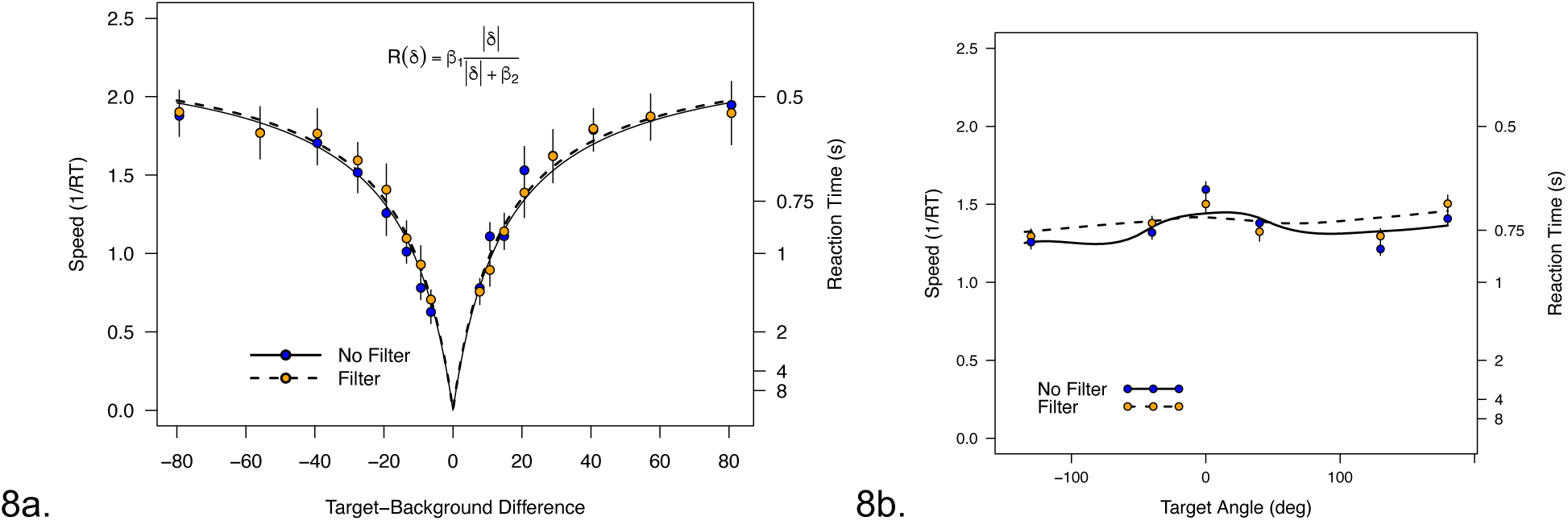
(a) Speed (1/RT) with and without the filter as a function of the TBD. Reaction times are indicated on the right ordinate. The points indicate the means with 95% confidence intervals computed across subjects and color coded the same as in Figure 7. The curves indicate a Michaelis-Menten function (inset equation) fit with a nonlinear mixed-effects model for the filter (dashed) and no filter (solid) conditions. Error bars correspond to 95% confidence intervals. (b) Speed as a function of the Target Angle (with reaction times indicated on the right ordinate). The points indicate the means and 95% confidence intervals computed across subjects. The curves are local regression fit (loess) to indicate the general trends through points.

We again fit the data with the NLME using Eq 2 to describe the dependence of the speed on TBD. As before we began by testing the significance of the random effect variance terms. As at –45°, the correlation between the two random effects for the model parameters was not significant (LRT: χ^2^(1) = 0.898, p = 0.34). Also, similarly to the –45° condition, both random effect variances were required to model the individual variability of the parameter estimates (LRT: 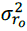 χ^2^(1) = 159.3, *p* << 0.001; 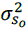, χ^2^ (1) = 5.52, *p* = 0.02). With the random effects structure established, we tested the fixed effects parameters. As for the –45° background, the estimate of the difference in values for maximum response, *R_f_*, did not differ significantly from 0 (LRT: χ^2^(1) = 1.862, p = 0.17) but the difference in values for the semi-saturation constant, ς*_f_* did (0 (LRT: χ^2^(1) = 7.53, p = 0.006). The parameter estimates for the fixed and random effects are indicated in column 4 Table 1 with their 95% confidence intervals in column 5. The fixed effect predictions of the two models are shown as solid and dashed curves in Fig 8a and suggest that while the filter effect on the semi-saturation constant is significant, it is very slight.

### Analyses of the search differences on the two backgrounds

Why was the filter enhancement on color search on the bluish-yellowish background more effective than on the SvsLM background? The basis for this difference is not readily evident in the context of the changes the filters induced in the cone opponent contrasts. We assumed that the salience of the targets was largely captured by the chromatic distance of the target from the background axis. As Figure 2 showed, filtering expanded chromatic contrasts along the magenta-lime direction. This left the –45*°* background axis largely unaffected, while rotating the 90*°* background by roughly 20°. However, the TBD remained similar across the two backgrounds, and in particular, the increase in distances with the filtering was also similar for the two conditions, as illustrated in Figure 9a. Moreover, while the 90*°* axis was rotated by filtering, the overall contrast remained similar, so that the chromatic variance along each background remained largely the same. Thus, based on the net cone-opponent contrasts, the effects of the filters on color search might be expected to be similar, in contrast to the differences we observed.

**Figure 9.**
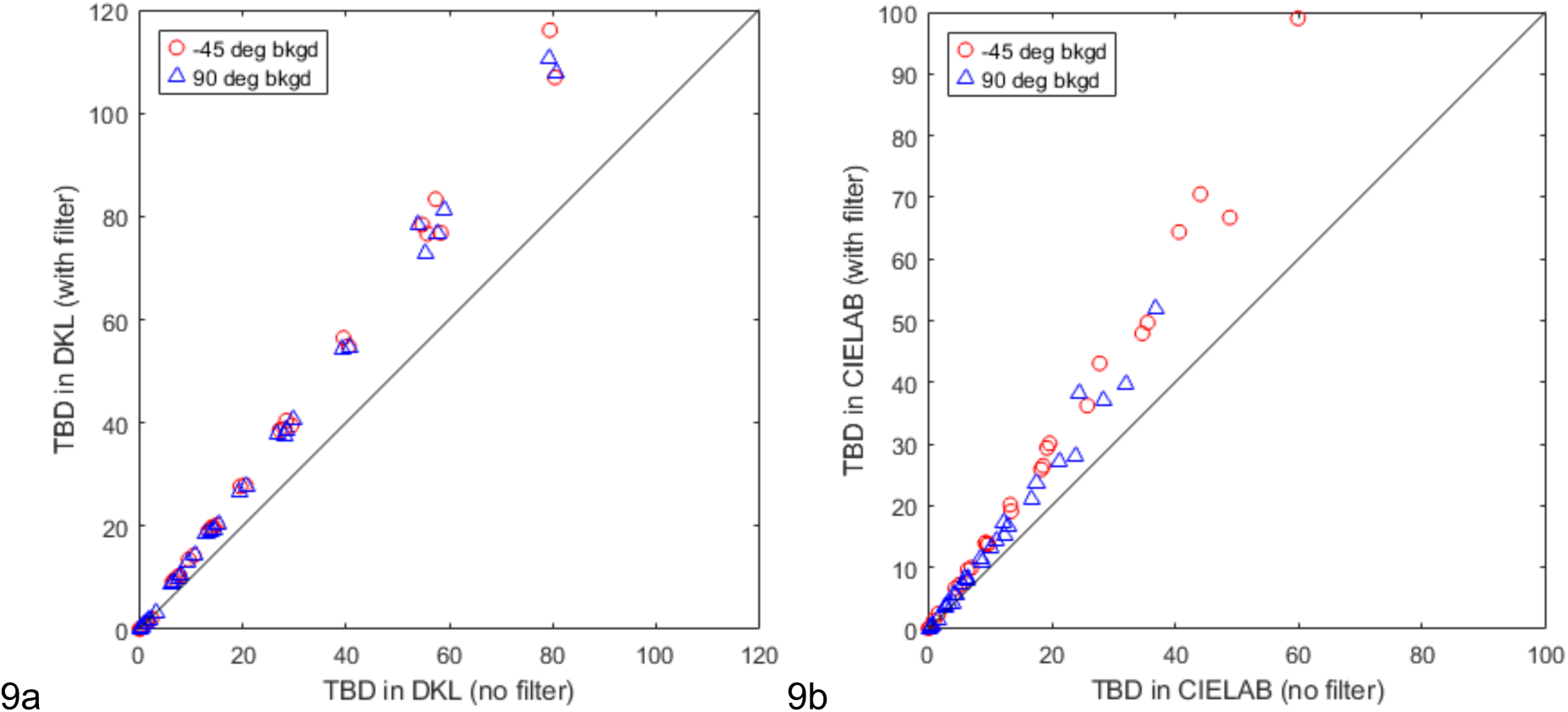
Chromatic target to background distance (TBD) for the –45° background (red circles) or the 90° background (blue triangles). a) TBDs within the cone-opponent plane. b) TBDs when the chromaticities are instead represented in CIELAB.

However, a clue to the basis for this difference is seen in average search times in Figures 5 and 6, which again show that locating targets on the –45*°* background was substantially faster than on the 90*°* background. This effect is consistent with previous results of McDermott et al. and Manalansan et al, who found that chromatic targets are easier to find on bluish-yellowish (or warm-cool) backgrounds than on orthogonal (magenta-greenish) backgrounds (Manalansan et al.,2025; McDermott & Webster, 2012). In turn, these effects are consistent with a large number of findings demonstrating weaker sensitivity to the bluish-yellowish axis of color space (Webster, 2020).

To explore this for our results, we converted the coordinates of the stimuli from the cone-opponent axes to the corresponding values in CIELAB, a “perceptually uniform” space designed to approximate perceptual rather than chromatic contrast differences. (Conversion to CIE DeltaE2000 yielded corresponding effects.) A circle in cone-opponent space, representing equal contrasts at different hue angles, warps into an elongated contour in LAB (McDermott & Webster, 2012). Manalansan et al. showed that the LAB contour minima correspond closely to the stimuli associated with warm and cool (orange and teal), while the maxima of the contours align with the categorical boundaries between warm and cool (Manalansan, Whitehead and Webster, 2025). In CIELAB, distances from gray approximate the chroma or perceptual strength of the color. The distortions thus imply that for the same cone-opponent contrasts, the perceptual strength is weakest along the warm-cool dimension and strongest along roughly orthogonal directions.

Figure 10 shows a similar analysis for our stimulus set and represents the LAB values for the target and background colors depicted in Figure 2 within the cone-opponent space. (Note that in addition to the warping there is an inversion of the blue-yellow axis between the spaces.) Within this perceptual metric, the 90*°* axis is expanded relative to the –45*°* axis. Moreover, the distances between the targets and background within the CIELAB space are markedly larger for the –45*°* background (Fig. 9b). We denote these values as TBD_lab_ to distinguish them from the TBD defined above for the cone-opponent plane. The asymmetries in the CIELAB values provide a plausible basis for the large asymmetries in search times on the two backgrounds.

**Figure 10.**
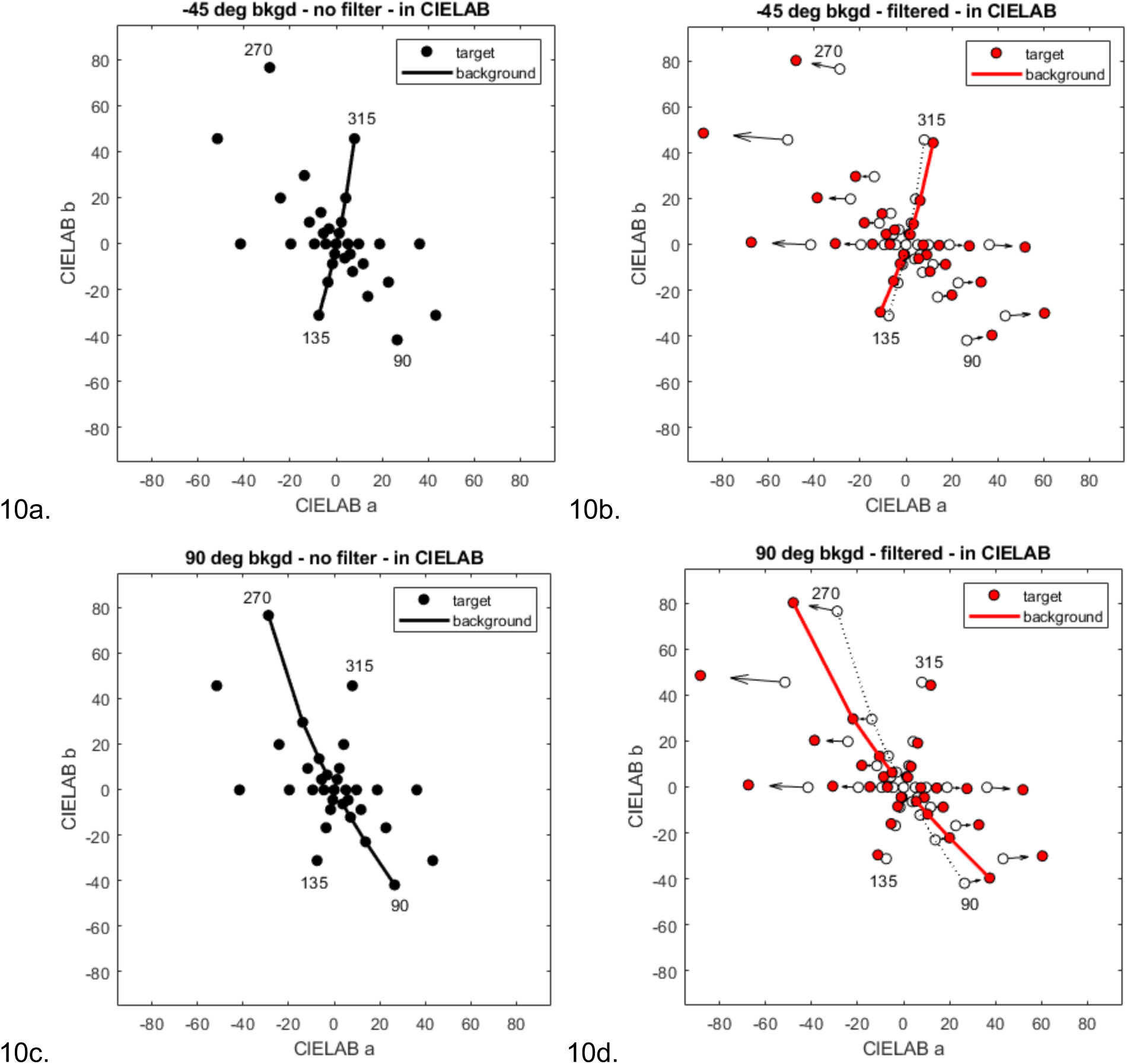
Chromaticities of the targets and backgrounds when converted to the the CIELAB color space. Unconnected circles show the coordinates of the targets, solid lines show the coordinates of the background axes. Labels indicate the original angles of the background axes in the cone-opponent space. a and b) Targets and –45 deg background before (a) or after (b) filtering. c and d) Targets and 90 deg background before (c) or after (d) filtering.

When the filter is applied, there is again little effect on the –45*°* background colors, while other stimuli are biased away from this axis. Thus, the faster search times with the filter are again consistent with the increased separation (TBD_lab_) of the targets from the background axis. In contrast, the filtering rotated the 90*°* background toward an axis of even greater perceptual sensitivity; and while the TBD_lab_ was also amplified, the enhancements relative to the background were weaker in the perceptually-based units. Thus, in this case the advantages of the filter for highlighting the targets are partially offset by the enhancement of the background. Finally, this analysis only considers the chromatic distance of the targets from the background axis, and not other factors such as the variance of the background. However, the contrast or range of the background colors also influences the salience of the targets, which for example are all much easier to detect on an achromatic background (McDermott et al., 2010). As noted, within the CIELAB matric this variance is higher for the 90*°* background, and modestly increased further by the filtering, and this is potentially an additional factor accounting for why filtering did not facilitate color search for this background for the specific conditions we tested.

## Discussion

To summarize, we found that the increases in chroma afforded by notch filtering led to predictable effects on color salience as measured by a color search task, and that the effects also depended in predictable ways on how the filter altered the chromatic properties of both the background and targets. In particular, for the conditions we tested, search times were significantly faster when the targets were shown on the bluish-yellowish background (–45*°*). In this case, the filter had only a marginal effect on the background chromaticities, while boosting other chromatic directions relative to the background axis, thus increasing the colorimetric distance from the background. On the other hand, for the purple to yellow-green background (90*°*), the targets were not easier to find even though the chromatic differences to the background were again increased. Notably, the basis for the lack of improvement on this background was only apparent when the stimuli were analyzed within a perceptual rather than cone-opponent space. While the backgrounds were equated in terms of cone-opponent contrasts, by the perceptual metric (CIELAB), the 90° deg background had a higher perceptual gamut than the blue-yellow background (Manalansan, Whitehead, et al., 2025; McDermott et al., 2010), and the filtering was less effective at selectively boosting the perceptual distances of the targets relative to this background.

Search times on the –45*°* background were also substantially faster than the 90*°* background, both with and without the filtering. This difference is again consistent with previous studies of color search using similar paradigms (Manalansan, Simoncelli, et al., 2025; McDermott et al., 2010). The weaker sensitivity and salience for the bluish-yellowish dimension has also been documented with a variety of other techniques e.g. (Bosten et al., 2015; Boynton et al., 1986; Goddard et al., 2010; Juricevic et al., 2010; Skelton et al., 2023). A common account of this bias is that the world varies more along the bluish-yellowish dimension, so that vision is more strongly adapted or desensitized to this axis (Juricevic & Webster, 2009; Skelton et al., 2024; Webster, 2020; Webster & Mollon, 1997). In this regard, adaptation also alters the relative salience of colors, but in a complementary way to the filtering we examined. Specifically, adaptation changes salience by reducing sensitivity to the background while “sparing” sensitivity to novel stimuli (McDermott et al., 2010; Wissig et al., 2013). In contrast, filtering is predicted to increase salience by enhancing the color of the target relative to the background (Somers & Bosten, 2024; Werner et al., 2020). In both cases, the changes in target salience should largely depend on how selective the effects are for the targets vs. the backgrounds. When the targets fall within the background distribution, salience may be relatively unaffected because both stimuli are biased in similar ways. Such predictions can also be readily extended to other background directions. For example, for the specific filter we examined, filtering the spectra should reduce color search on LvsM (e.g., 0-180°) backgrounds, where the effect would instead be to enhance the background colors relative to the targets. However, such color backgrounds are atypical, at least for natural environments.

Could differences in adaptation account for the effects we observed for the simulated filters? There are at least two principal types of this adaptation (Webster, 1995). The first is chromatic adaptation to the average color and luminance of the display. As noted in the Methods, we corrected the stimuli for this factor by rescaling the predicted cone responses so that the mean responses were the same before or after filtering. As a result, all conditions had the same average background and thus should have the same average state of chromatic adaptation.

The second form of adaptation is to the contrast or variance of the colors. Observers selectively adapt to the angle and variance of color distributions (Krauskopf, Williams, and Heeley, 1982; Webster and Mollon, 1994), and in previous work using very similar stimuli and tasks we showed that this adaptation enhances the salience of target colors that differ from the background (again as inferred from shorter search times for the targets) (McDermott et al., 2010). In the present study we did not attempt to control for differences in contrast adaptation for the filtered vs. unfiltered conditions, though effects of long-term exposure have been found in previous work (Werner et al., 2020; Somers and Bosten, 2024). However, contrast adaptation is unlikely to account for the differences we observed across the present conditions. In particular, as Figure 2 shows, the filtering had only a small effect on the angle or range of the colors for the – 45 deg background. The backgrounds colors should set the state of adaption, since the target colors are in comparison very rare. Thus for this background no differential adaptation would be expected between the filter and no filter conditions, in contrast to the effects we observed.

As noted in the Introduction, the two backgrounds we examined corresponded to the S-cone modulating or tritan cardinal axis of cone opponency (900) vs. an axis closer (but not identical) to the blue-yellow principal axis of perceptual opponency. These were chosen because theyroughly span the range of dominant chromatic axes of many natural scenes – and in particular capture the color signatures of lush or arid environments (Nascimento et al., 2021; Ruderman et al., 1998; Skelton et al., 2024; Webster & Mollon, 1997). However, in other ways these backgrounds were also highly unnatural: the colors were confined to a single hue angle and sampled the axis uniformly, and had high variance. In contrast, natural color distributions are more irregular and clustered (e.g., between regions of sky and earth), and often have a lower variance with higher proportion of low-contrast levels (Laughlin, 1981; Nascimento et al., 2002; von der Twer & MacLeod, 2001). Moreover, in real scenes colors are usually associated with objects and surfaces and thus have very different spatial characteristics from the uniform textures we used. A further limitation is again that we simulated the filters rather than directly wearing them, and did not evaluate the effects the filters have on the mean light level (Somers & Bosten, 2024). The simulations did, however, capture the pattern of cone excitations predicted if observers in each case were adapted to the mean level of the display. This light and chromatic adaptation should largely offset the effects of absolute intensity differences but does not fully address potential changes in chromatic discrimination at different light levels.

The highly constrained conditions we tested are useful for controlling the stimuli and revealing the principles affecting the search performance, but are unlikely to fully generalize to realistic viewing environments. As one example, we used the target to background distance (TBD) as a simple predictor for salience. In real scenes, however, the background distribution will not be confined to a single axis, so that the target salience is more likely to be a function of the signal to noise ratio. By this metric, even targets that lie along the dominant axis can (and do) become salient as they fall outside the background gamut (McDermott et al., 2010; Manalansan, Simoncelli, et al., 2025). In such cases, an important question for understanding the effect of the filters is how amplifying both background and target (while for example maintaining the contrast ratio) might impact the color search. Similarly, in natural scenes, search is strongly affected by the specific task demands and expectations about the potential presence, location, or nature of specific targets, and thus factors such as expectations and familiarity, both with objects and their colors, will also impact the salience (Võ, 2021; Wolfe & Horowitz, 2017). In future work we plan to evaluate the contrast enhancements under these more naturalistic conditions.

A further factor determining salience is how the color signals are represented in the visual system (at least at the level limiting performance). For example, as our results revealed, the different effects of the filtering could not be explained by the cone-opponent contrasts in the stimuli, and instead depended on sensitivity biases that were better captured by a color space that approximates perceptual uniformity. This bias itself argues against stages of color coding where the LvsM and SvsLM cardinal axes are independent, because the blue-yellow bias (i.e. at –45*°*) occurs relative to stimuli that are matched for the cardinal axis responses (i.e. for axes at +45*°*, where the LvsM and SvsLM signals are equal but combined in opposite phase) (Goddard et al., 2010). Finally, predicting the salience also depends on the underlying structure of the color coding. Our TBD metric assumed that backgrounds had no effect on orthogonal target directions. However, at cortical stages where color is encoded by multiple hue selective mechanisms (Gegenfurtner, 2003), these orthogonal signals can interact. For example, the broad color tuning at early cortical levels means that while cells tuned to 0*°* or 90*°* do not “see” both axes, cells tuned to intermediate axes (e.g. 45*°*) respond and contribute to both (Webster & Mollon, 1994). Thus, it is an oversimplification to assume that there is no effect of the backgrounds on the orthogonal targets. Finally, we have considered only the changes in chromatic contrast and perceived hue and chroma. However, increases in saturation are also associated with changes in perceived brightness, which could serve as a further potential cue for salience (Corney et al., 2009). A more formal model of color search would need to incorporate these different factors.

Our results have a number of implications for the potential to enhance color vision through spectral filtering. First, these results are specific to the specific filter design we evaluated, which again was developed for normal trichromatic vision. In other designs the notch filtering has been tailored to enhance chromatic contrasts for different types of anomalous trichromacy (i.e., deuteranomalous or protanomalous) (Somers & Bosten, 2024). The current study tested only normal color vision, but the color search task could be readily adopted for assessing the impact of the filters for color deficiencies. An advantage of the paradigm is that it can evaluate a wide range of both threshold and suprathreshold colors while exploiting a common and naturalistic visual task. The present analyses and measurements are also relevant to understanding consequences of varying the illumination rather than filtering. As noted, wide gamut lighting with narrowband primaries applies similar principles to enhance chromatic contrasts, and thus predicts similar consequences for color salience (Ilic et al., 2022), for both normal and color-anomalous vision (Tamura, Okamoto et al., 2017ì5; Tamura, Okamoto et al., 2017; Joyce et al., 2021). Finally, filters (or lighting) could also be engineered for different environments or goals, as well as for different observers, depending on the availability of appropriate dyes. For example, filters can be designed to minimize the changes specifically along the 90*°* axis rather than the bluish-yellow axis of the design we tested here. This selectively expands chromatic contrasts relative to the SvsLM opponent axis and thus better isolates the signals in the LvsM axis, and is the principle used in designs for color vision deficiencies. The LvsM signals are carried by the “modern subsystem” of primate trichromacy and may have evolved specifically to make ripening fruit more distinguishable from the foliage background, by restricting the chromatic noise from the background of foliage to primarily SvsLM variations (Osorio & Vorobyev, 1996; Regan et al., 2001; Sumner & Mollon, 2000). As the two backgrounds we tested again illustrate, natural (as well as carpentered) environments do vary substantially across both location and time (e.g., seasons (Webster et al., 2007)). Adaptation is thought to help tune vision for specific environmental contexts (Webster, 2015). In a similar way, notch filtering could potentially be optimized for specific environments or tasks.

## Acknowledgments

Supported by DOD W911NF-24-1-0328 (DM, MW, KK), NIH EY010834 (MW) and LABEX CORTEX (ANR-11-LABX-0042) of the Université de Lyon (ANR-11-IDEX-0007) from the French National Research Agency (KK).

